# A Cortical Site that Encodes Vocal Expression and Reception

**DOI:** 10.1101/2024.10.15.618282

**Authors:** Thomas Pomberger, Katherine S Kaplan, Rene Carter, Thomas C Harmon, Richard Mooney

## Abstract

Socially effective vocal communication requires brain regions that encode expressive and receptive aspects of vocal communication in a social context-dependent manner. Here, we combined a novel behavioral assay with microendoscopy to interrogate neuronal activity in the posterior insula (pIns) in socially interacting mice as they switched rapidly between states of vocal expression and reception. We found that distinct but spatially intermingled subsets of pIns neurons were active during vocal expression and reception. Notably, pIns activity during vocal expression increased prior to vocal onset and was also detected in congenitally deaf mice, pointing to a motor signal. Furthermore, receptive pIns activity depended strongly on social cues, including female odorants. Lastly, tracing experiments reveal that deep layer neurons in the pIns directly bridge the auditory thalamus to a midbrain vocal gating region. Therefore, the pIns is a site that encodes vocal expression and reception in a manner that depends on social context.

## INTRODUCTION

Vocal communication is an essential medium for forging and maintaining social bonds in all mammalian species, including humans^1–4^. Socially effective vocal communication requires that vocal expression and reception (i.e., listening) are carefully regulated as a function of social context. For example, vocalizations require an audience to exert their social effects, and the audience must in turn discern which vocalizations signify socially relevant exchanges. A major unresolved issue is the extent to which single brain regions encode expressive and receptive aspects of vocal communication in a manner that is sensitive to social context.

The insular cortex binds various sensory and social signals to guide behavior^5–11^, providing a potential site for encoding socially salient vocal signals. In fact, the posterior insula (pIns) integrates multisensory information, including auditory stimuli, in monkeys^12^ and mice^13–15^. In monkeys, pIns neurons respond to a range of animal vocalizations, with the strongest responses evoked by conspecific vocalizations^12^. Although pIns activity during vocal expression has yet to be described in monkeys or rodents, intracranial electroencephalography (iEEG) recordings in human subjects show enhanced activity during speech as well as speech playback^16^, and human patients with insula lesions suffer from articulatory planning deficits^17,18^. Therefore, the pIns is an attractive candidate brain region where expressive and receptive aspects of vocal communication may be encoded in a manner that is sensitive to social context.

Our understanding of how the pIns encodes expressive and receptive aspects of vocal communication is currently limited. First, most studies of the auditory properties of pIns neurons have presented vocalizations through a speaker to head-fixed, socially isolated animals, a state where vocalizations are devoid of social context. Furthermore, a systematic characterization of the same populations of pIns neurons during social interactions that involve both vocal expression and reception have yet to be undertaken. While the recent characterization of pIns in humans is a step in this direction, iEEG lacks cellular resolution and how social context modulates pIns activity remains unknown.

Here, we combined a novel behavioral assay with miniature microendoscopy (miniscope) in which we could interrogate pIns neuronal activity in socially interacting mice as they switched rapidly between states of vocal expression and reception. We found that distinct but spatially intermingled subsets of pIns neurons were active during these two states. Notably, pIns activity during vocal expression increased prior to vocal onset and was also detected in congenitally deaf mice, consistent with a motor-related signal. Moreover, the subset of pIns neurons that were activated when a mouse listened to vocalizations produced during social encounters were activated only weakly or not at all by vocal playback when the mouse was by itself. Further analysis of pIns activity using multiphoton imaging in head-fixed male mice in which we could carefully regulate exposure to a female mouse revealed that female odorants enhanced pIns responses to vocal playback. Lastly, tracing experiments reveal that deep layer neurons in the pIns directly bridge the auditory thalamus to a vocal gating region in the periaqueductal gray (PAG). These findings identify the pIns as a site where auditory and motor representations of vocal communication signals are represented in a manner that depends on social context.

## RESULTS

### A behavioral protocol for monitoring social-vocal communication

Male mice emit ultrasonic vocalizations (USVs) when exposed to female mice or their odors^19–22^, and these vocalizations facilitate mating^23,24^. This courtship behavior provides an ethologically relevant context in which to explore the neural correlates of expressive (in the male) and receptive (in the female) aspects of social communication. Furthermore, while the female mouse is the intended audience of the male’s USVs, in natural settings other mice including rival males can eavesdrop on these vocal bouts and thus detect these courtship encounters. Therefore, eavesdropping males provide an additional context in which to probe the neural correlates of vocal reception.

In order to study the neural correlates of these expressive and receptive processes, we developed a two-chamber system in which to probe neural activity in the pIns in male and female mice during social encounters in which the males typically emit USVs (Fig. 1A). In this setup, a male mouse was housed with a female in a “courtship” chamber while another male was placed in an adjacent “eavesdropping” chamber. The two chambers were separated by a mesh screen through which auditory signals and odors were transmitted. The movements of the individual mice were monitored under infrared illumination, eliminating any social signals provided by visual cues. Microphones over each chamber were used to detect USVs and to establish that vocalizations emanated exclusively from the courtship chamber. We assumed that the majority of these USVs were produced by the male in the courtship chamber, given that female mice rarely emit USVs during courtship encounters^25–27^. Therefore, both the female as well as the male in the adjacent chamber served as receivers for the courting male’s USVs. By moving the female from one chamber to the other, we were able to switch a male’s role from emitting USVs during courtship to eavesdropping on the other male’s courtship USVs. Finally, we isolated the experimental mouse and delivered a series of pre-recorded male USVs through a speaker, allowing us to measure auditory responses to vocalizations in the absence of social cues.

**Figure 1:**
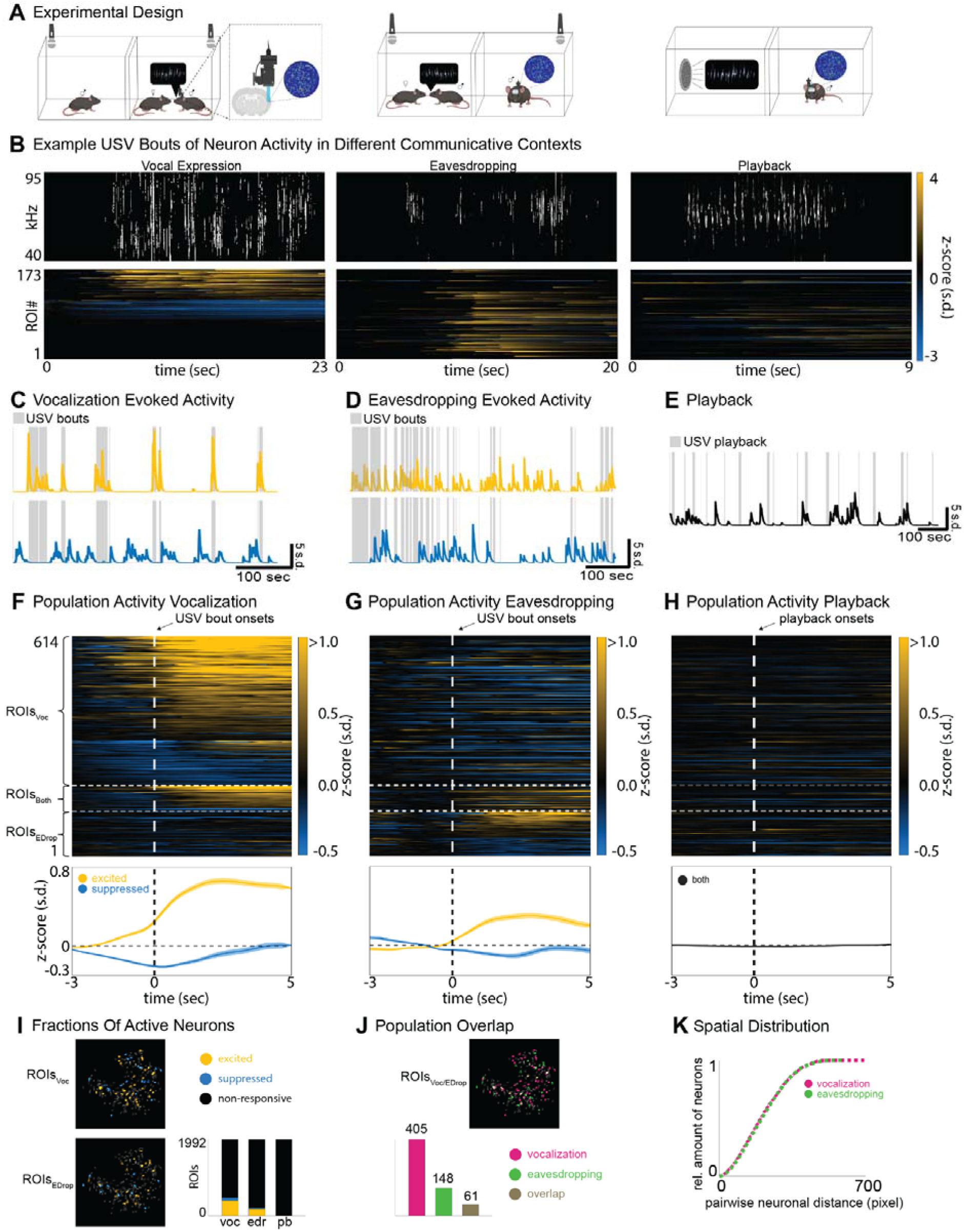
The posterior insula encodes expressive and receptive aspects of social-vocal communication. (A) Experimental design showing the three different social-vocal contexts. (B) Example USVs during vocal expression (top left), eavesdropping (top middle) and playback (top right), and the corresponding ROI activities below. (C-E) Example ROIs showing activity during USV bouts and USV playback for each of the three communicative contexts (yellow = excited, blue = suppressed, black = non-responsive). (F-G) Average activity of ROIs that are active during vocal expression (ROIs_Voc_), during eavesdropping (ROIs_EDrop_) or in both contexts (ROIs_Both_). Top panels show each individual ROI. The bottom panels show the overall population activity of excited (yellow) and suppressed (blue) ROIs. (H) Same ROIs as in G & H but shown during USV playback. (I) Example field of views of ROIs_Voc_ and ROIs_EDrop_ (top and bottom left) and amount of excited, suppressed and non-responsive ROIs in each context (voc = vocal expression, edr = eavesdropping, pb = playback). (J) Example field of view of ROIs that are responsive to vocal expression or eavesdropping and their overlap (top). Total amount of responsive ROIs in vocal expression (magenta), eavesdropping (green) or both contexts (brown). (K) Cumulative distribution function of pairwise neuronal distances in pixel of ROIs_Voc_ (magenta) and ROIs_EDrop_ (green).

### The posterior insula is active during socially salient vocal expression and reception

We combined our behavioral approach with calcium imaging using a miniature microscope (miniscope) to monitor pIns activity during expressive and receptive phases of socially salient vocal communication and when vocal stimuli were presented in social isolation. Briefly, we used viral vectors to express GCaMP8s pan-neuronally in the pIns and a GRIN lens to gain optical access to superficial and deep layers of this region (Fig. 1A, left). Qualitatively, activity of pIns increased sharply in male mice when they emitted USVs during courtship interactions with a female and when they eavesdropped on live USV bouts of another male suitor (Fig. 1B, left and middle). In contrast, pIns neurons were only weakly activated in trials where socially isolated males listened to USV bouts played through a speaker (Fig. 1B, right).

To more systematically quantify these effects, we performed a receiver operating characteristic (ROC) analysis, allowing us to identify subsets of neurons that were significantly excited or suppressed relative to baseline during USV production, eavesdropping and playback (Fig. S1A). This approach confirmed that subsets of ROIs in the pIns were significantly excited or suppressed during vocal production and eavesdropping (Fig. 1C and fig. 1D, left top and bottom), but not during playback (Fig. 1E). Furthermore, the ROC analysis revealed that the population of pIns ROIs active during self-produced vocalizations (pIns_Voc_) was mostly non-overlapping with the population active during eavesdropping (pIns_EDrop_) (Fig. 1F and fig. 1G, right top and bottom). A smaller number of ROIs were modulated during self-produced vocalizations and eavesdropping (Fig. 1F and fig. 1G, right middle). In contrast, pIns neurons were barely activated when socially isolated mice listened to the same vocalizations played through a speaker (Fig. 1H). In total, ∼22% (466 of 1992) of ROIs in the pIns were significantly modulated from baseline during self-initiated vocalizations and ∼10% (209 of 1992) were modulated during eavesdropping (Fig. 1I, N = 5 male mice, Wilcoxon, p < 0.05). Furthermore, ∼9% of all responsive ROIs (61 of 675) were significantly modulated from baseline during both vocal production and eavesdropping (Fig. 1J). In contrast, only 1 ROI was significantly modulated by USV playback. Therefore, different populations of neurons in the pIns encode expressive and receptive aspects of vocal signals and auditory responses to these signals are sensitive to the social context in which they are heard.

A notable feature of the population of pIns_Voc_ neurons was that their activity deviated from baseline prior to vocal onset, whereas activity in the pIns_EDrop_ population deviated after the onset of the other male’s USVs (Fig. 1F and fig. 1G, bottom). Therefore, modulation of pIns activity during vocal production was not purely a consequence of vocalization-related auditory feedback and instead may reflect a premotor signal. We also examined whether these two populations of pIns neurons were spatially distinct. Qualitatively, pIns_Voc_ and pIns_EDrop_ appeared to be intermingled across the imaging field of view (Fig. 1I & 1J, top). Furthermore, the probability of pairwise Euclidean distances between pIns_Voc_ and pIns_EDrop_ ROIs were closely overlapping, indicating these two populations have similar spatial distributions in the insula (Fig. 1K, two-sided KS test, p = 0.71). In summary, largely distinct populations of spatially intermingled pIns neurons are active during vocal expression and reception in male mice, and may separately encode motor versus auditory information about USVs.

### Activity during vocal expression is not attributable to locomotion

Vocalization in freely behaving male mice typically occurs during female pursuit, and locomotion can modulate activity in sensory cortices, including the auditory cortex^28–30^. In fact, we confirmed that the male’s running speed increased prior to USV onset, raising the potential confound that vocal modulation of pIns activity was driven by locomotion rather than vocal production (Fig. 2A). However, aligning pIns_Voc_ ROIs to either running onset or acceleration in running speed failed to detect any change in fluorescence (Fig. 2B & 2C, Wilcoxon, p = 0.31). Furthermore, we trained a long short-term memory (LSTM) network to decode either vocalization or running from the entire population of pIns ROIs (1992 ROIs from N = 5 male mice). The decoding accuracy of the LSTM was significantly greater for vocalization when compared to shuffled data, while running state could not be decoded (Fig. 2D & 2E, Wilcoxon ranksum, p < 0.05 & p = 0.69). Therefore, activity in pIns_Voc_ ROIs is a consequence of vocal production rather than locomotion.

**Figure 2:**
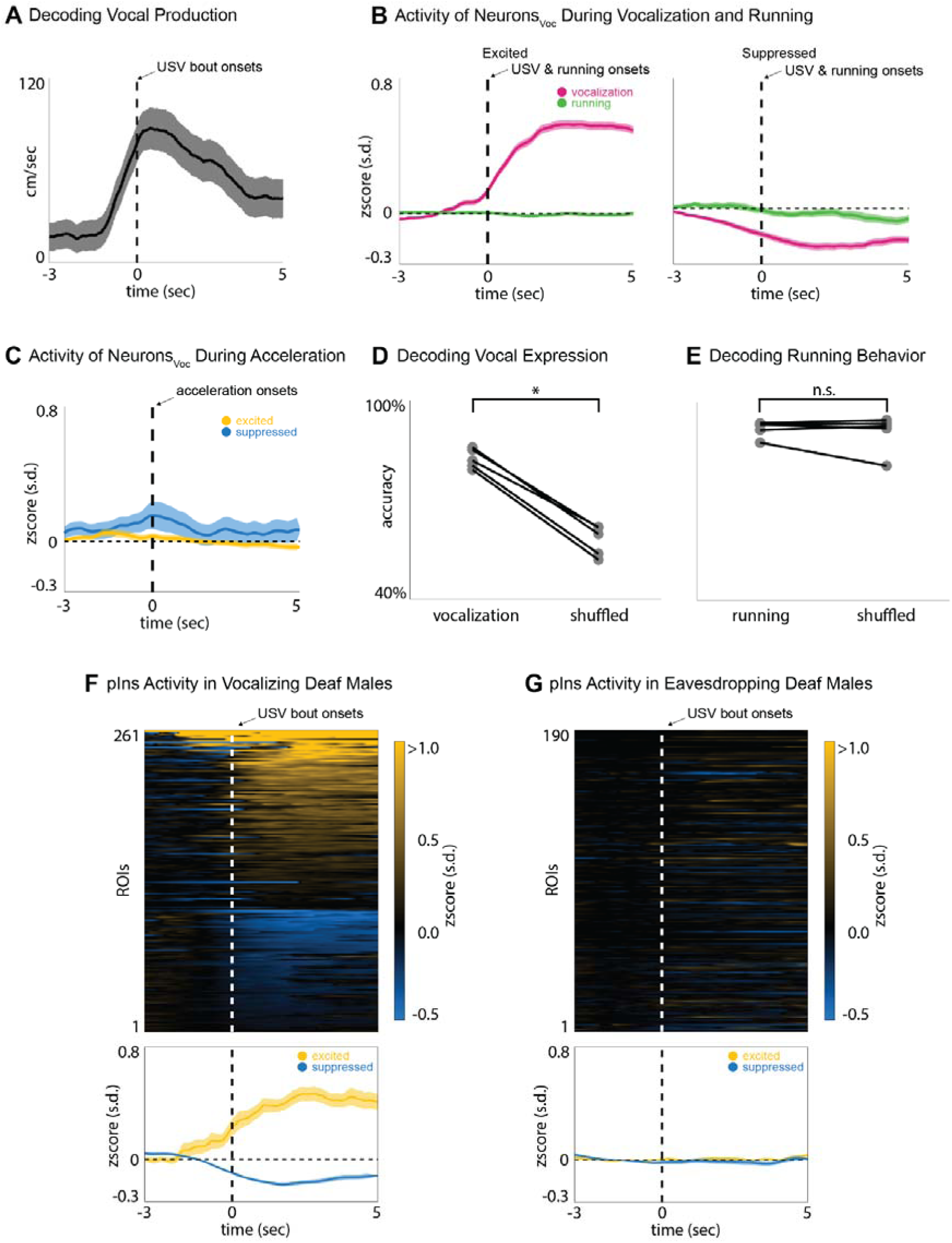
Activity in the posterior insula encodes vocal expression in hearing and deaf mice. (A) Average running speed of vocalizing male aligned to USV bout onset. (B) Average population activity of excited (left) and suppressed (right) ROIs_Voc_ aligned to USV bout onset (magenta) and running bout onset (green). (C) Average population activity of excited (yellow) and suppressed (blue) ROIs_Voc_ aligned to acceleration bout onset. (D-E) Decoding accuracies for vocal expression (right) and running (left). (F) Average activity of ROIs that are active during vocal expression in deaf males (top) and the corresponding average population activity of excited (yellow) and suppressed (blue) ROIs. (G) Same is in (F) but for activity during eavesdropping in deaf males.

### Activity during vocal expression does not require hearing

As previously noted, activity in pIns_Voc_ ROIs increased prior to vocal onset, indicating that activity during vocal expression is not limited to vocalization-related auditory feedback. To further probe the extent to which vocal modulation in pIns_Voc_ ROIs was independent of auditory feedback, we monitored pIns activity in congenitally deaf males as they engaged in courtship or eavesdropping (Tmc1(Δ))^31^. Calcium signals in a subset of pIns ROIs in Tmc1(Δ) male mice were modulated from baseline during vocal expression (Fig. 2F). Notably, the proportion of vocalization-modulated pIns_Voc_ ROIs (∼25%, 261 of 1063 ROIs, N = 5 males) and the time course of their vocal modulation was similar in deaf and hearing mice. Therefore, modulation of pIns activity during vocal expression does not require auditory feedback, pointing to the presence of either a motor or a proprioceptive signal. In contrast, no modulation occurred when Tmc1(Δ) males were placed in the eavesdropping chamber and exposed to another male’s courtship USVs (Fig. 2G, N = 4 males), confirming that eavesdropping-related activity depends on hearing.

### pIns Neurons in the female mouse respond to socially salient USVs

We also imaged pIns neurons in female mice housed in the courtship chamber with a vocalizing male (Fig. 3A). A subset of pIns ROIs in females responded strongly to USVs of a male suitor (Fig. 3C & 3D). An ROC analysis quantified 153 of 1500 ROIs as USV-responsive, similar to the proportion of USV-responsive pIns ROIs detected in eavesdropping males (Fig. 3E, N = 5 females). As in eavesdropping males, the responses of female pINs ROIs to male USVs were strongly dependent on social context, as no ROIs were modulated by USV playback when these females were in social isolation. In summary, a subset of pIns ROIs are active during vocal reception in both female and male mice.

**Figure 3:**
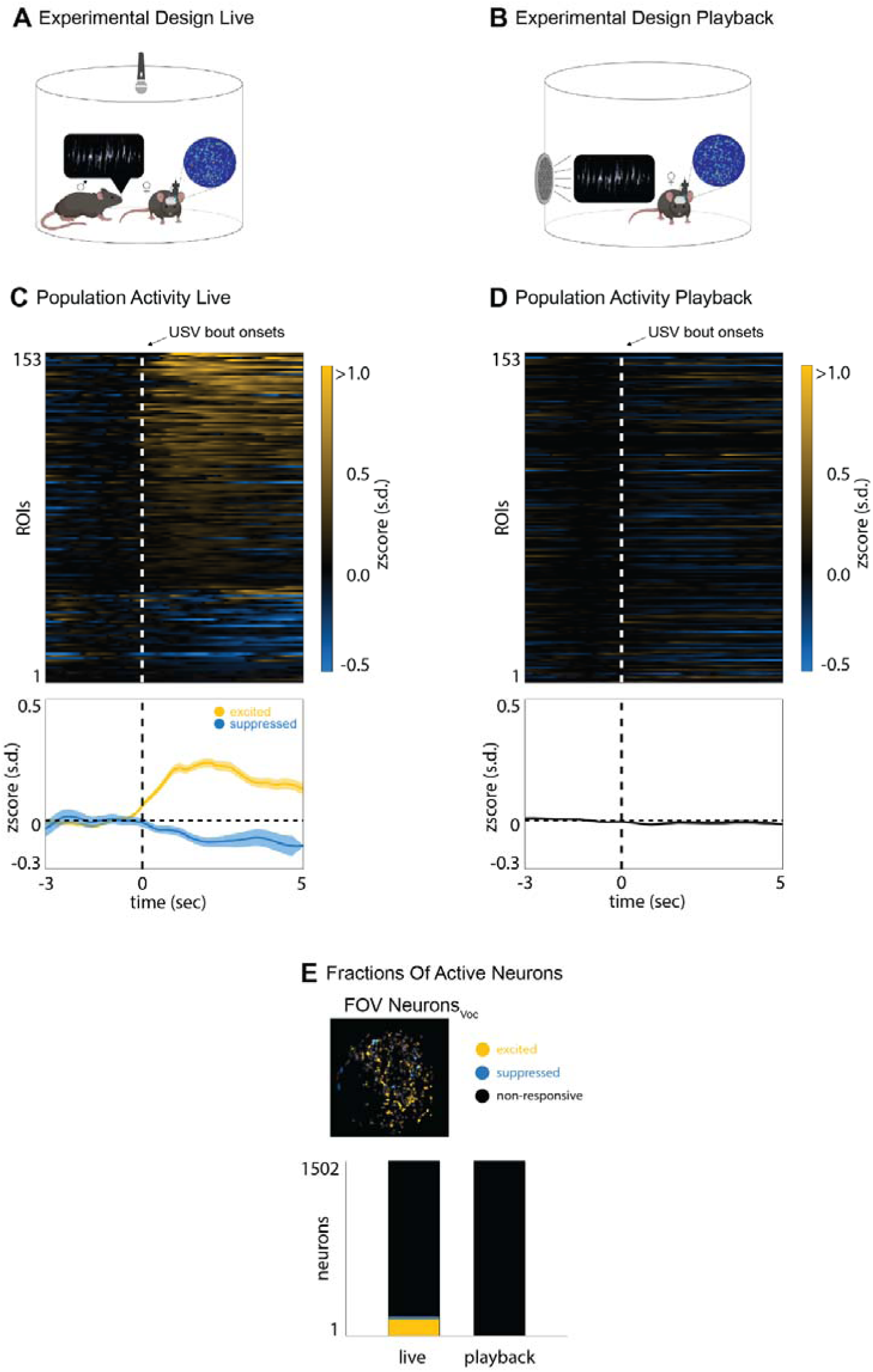
The posterior insula of females responds to social vocalizations of males. (A-B) Experimental design for live and playback vocalizations. (C) Average ROI activity in the posterior insula of females while exposed to a vocalizing male (top). The bottom panel shows the average population activity of excited (yellow) and suppressed (blue) ROIs. (D) Same ROIs as in (C) but shown during USV playback. (E) Example field of view of excited (yellow) and suppressed (blue) ROIs of a female when exposed to a vocalizing male (top). Total amount of ROIs for each of the two contexts (live vocalization and playback).

### Female Odor Increases Auditory Responsiveness in Male Posterior Insula

Here we found that a subset of neurons in the pIns of eavesdropping male mice respond strongly to USVs produced by a nearby courting male, whereas USV playback elicits only weak responses in the pIns when males are in social isolation. Therefore, additional non-vocal social cues must augment responses of pIns neurons to the other male’s USVs. Given the multi-sensory nature of pIns^14,32^, we hypothesized that female odor is one of these social cues. Regulating odor delivery in unrestrained courting mice is impractical, so we instead used 2p methods to image pIns activity in the head-fixed male mouse while regulating its exposure to female mouse odors and delivering pre-recorded USVs of other males through a speaker. In this setting, odors were delivered to the head-fixed male by directing airflow into a chamber containing the female and through a nozzle in front of the male’s snout (Fig. 4A, middle). Under conditions of no directed airflow or when the female was absent (Fig. 4A, top), a subset of pIns ROIs were either excited or suppressed by USV playback (Fig. 4B, top and bottom; the proportion of playback responsive ROIs was ∼37%, 499 of 1351 ROIs; this was significantly higher than USV playback-excited neurons detected using miniscopes in socially isolated male mice; Wilcoxon ranksum, p < 0.05). When airflow was directed towards the male, the magnitude of the excitatory and suppressive responses of these ROIs to USV playback increased (Fig. 4B, middle and bottom, fig. S2A, N = 4 males, Wilcoxon, p < 0.05). Separate from playback responses, we did not detect any differences in fluorescence between the undirected (i.e., no odor) and directed airflow conditions (Fig. S2B, two-sided KS test, p = 0.17). Therefore, female odorants modulate auditory responses in the male’s pIns to other male’s USVs.

**Figure 4:**
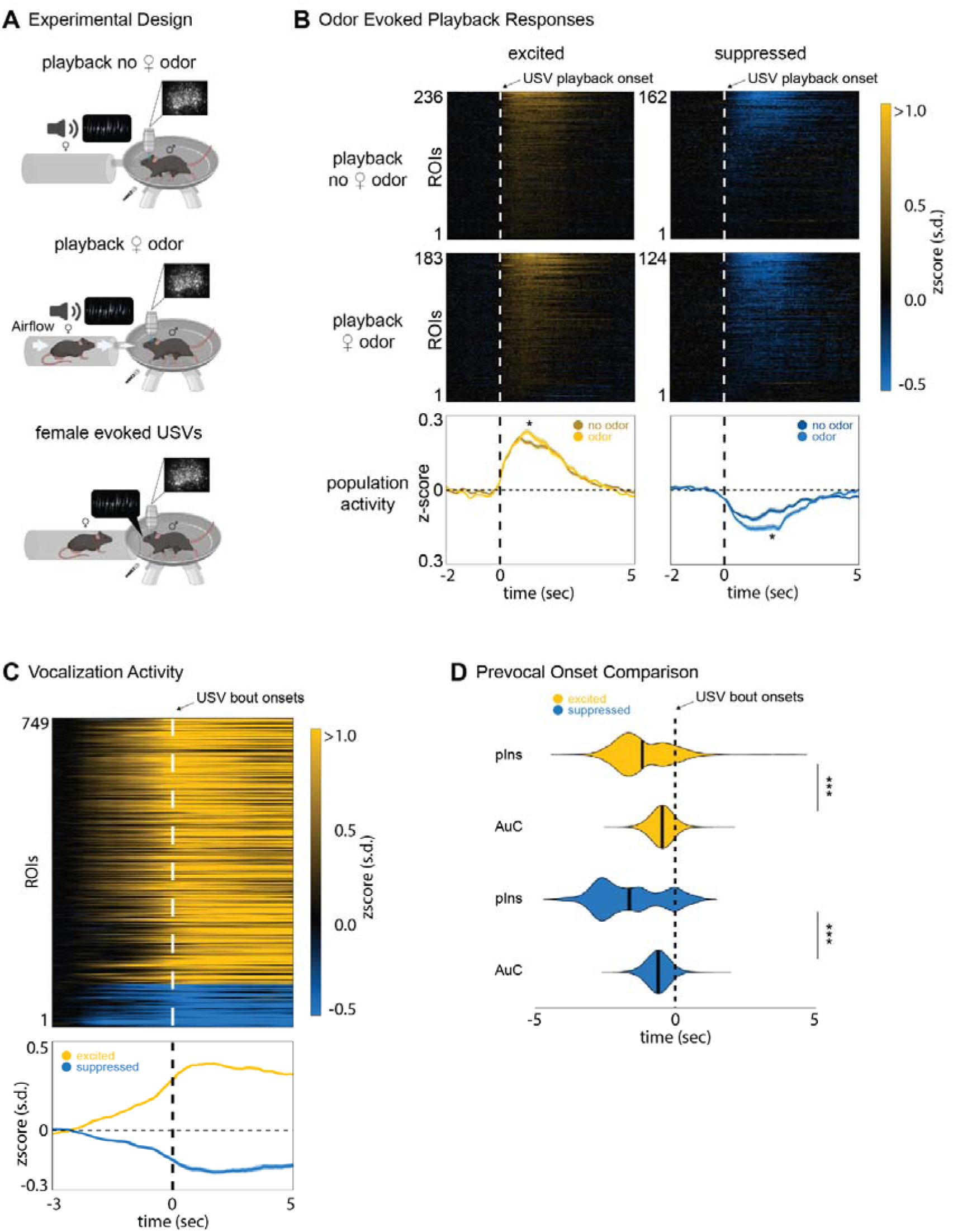
Female odor increases responsiveness in the posterior insula of males. (A) Experimental design showing the head-fixed male exposed to USV playback during neutral airflow (top), positive airflow that delivers odorants from a distal female (middle) and USVs elicited by an approaching female (bottom). (B) Average activity of ROIs that were active during neutral and positive airflow (top and middle). The bottom panel shows the average population activity of excited (yellow) and suppressed (blue) ROIs during neutral and positive airflow. Stars indicate significance between the two populations. (C) Average activity of ROIs that were active during vocal expression in head-fixed males. Bottom panel shows average population activity of excited (yellow) and suppressed (blue) ROIs. (D) Prevocal onset activity of posterior insula (pIns) and auditory cortex (AuC) of excited and suppressed ROI populations.

We also conducted an additional experiment in which a female mouse could approach the head-fixed male mouse snout to snout, which often elicited USVs from the male (Fig. 4A bottom). As in the miniscope experiments, pIns activity was strongly modulated in the pIns of vocalizing, head-fixed males, and this vocalization-related activity increased before vocal onset (Fig. 4C). In fact, 2p imaging detected an even greater proportion of pIns ROIs that were modulated during USV production (∼55%, 749 of 1351 ROIs). However, the subset of pIns ROIs that were modulated during vocalization was largely non-overlapping with those that were excited or suppressed by USV playback in either the presence or absence of female odor (∼33% overlap of all responsive ROIs). Finally, we compared the activation times of these vocalization modulated ROIs in pIns with those of vocalization modulated ROIs in auditory cortex (AuC) from a previous study^30^. This comparison revealed that the modulation prior to vocal onset occurred earlier in the pIns than in the AuC (Fig. 4D, Wilcoxon, p < 0.001). In summary, the 2p imaging approach used here revealed that a greater proportion of pIns ROIs were active during USV production and playback than detected using miniscopes, presumably reflecting the enhanced sensitivity of 2p imaging methods. Nonetheless, 2p imaging confirmed that the subsets of ROIs activated during these two conditions were largely non-overlapping while also showing that female odorants modulate male pIns activity evoked by auditory presentation of other males’ USVs.

### The Posterior Insula Links the Auditory Thalamus with a Vocal Gating Region in the PAG

Previous neuronal tracing studies with fluorescent markers provide evidence that neurons in the auditory thalamus (MGB) make axonal projections to the pIns^13–15^. To further characterize this projection, we injected retrograde AAV-Cre (AAVrg-Cre) in the pIns and AAV-flex-GFP in the MGB (Fig. 5A left, N = 3). This approach resulted in robust GFP expression in cell bodies in the medioventral part of MGB (MGB_pIns_) and in the posterior intralaminar thalamic nucleus (PIL), which is adjacent to the MGB and is implicated in social, maternal and sexual behaviors^33–36^ (Fig. 5A right). This intersectional approach also resulted in dense GFP labeling in axon terminals in all layers of the pIns (Fig. 5A middle), confirming that thalamic regions including the MGB and PIL are a major source of input to the posterior insula.

**Figure 5:**
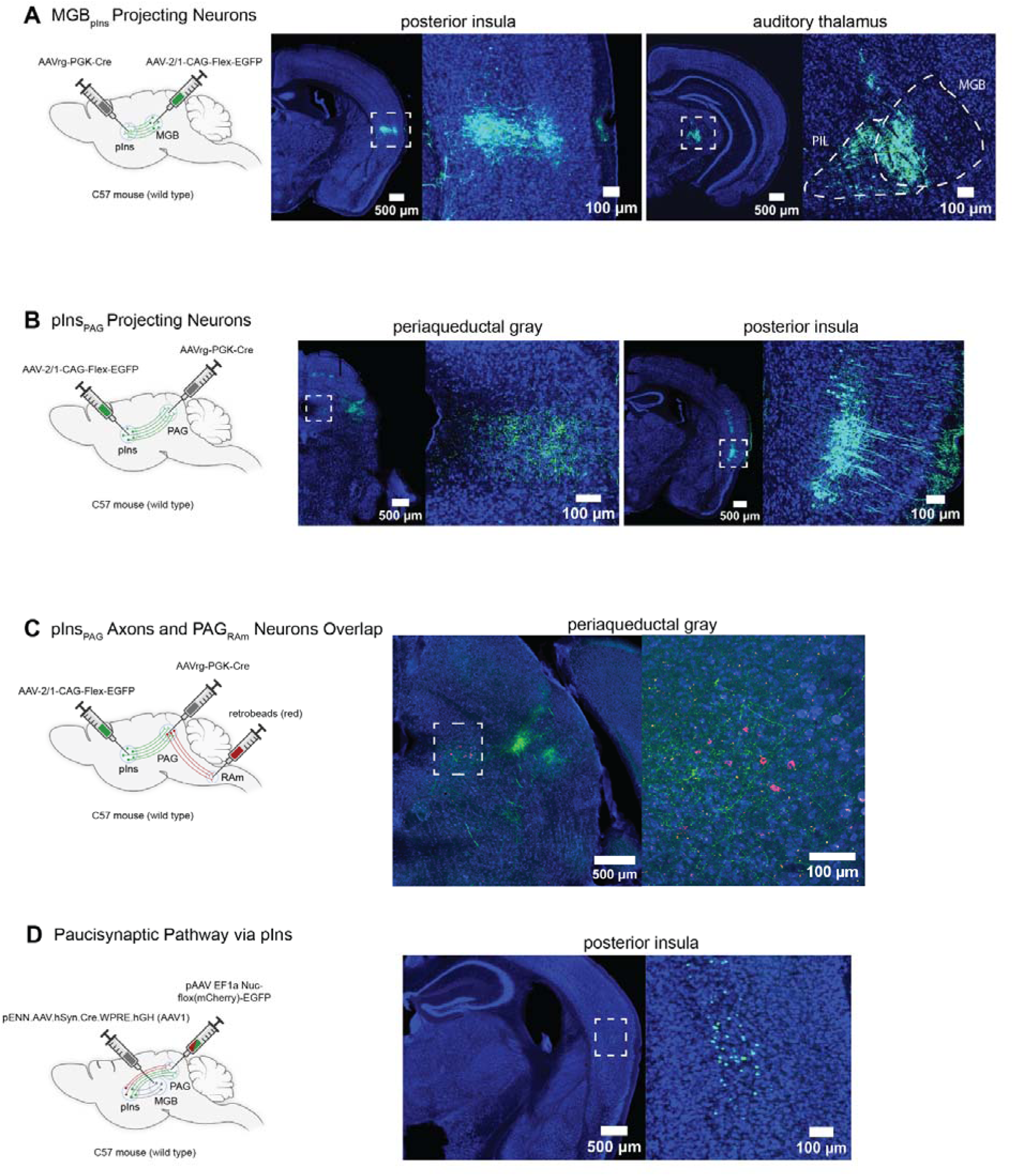
The posterior insula links the auditory thalamus with a vocal gating region in the PAG. (A) Experimental design to label MGB_pIns_ projecting neurons (left); axon terminals in pIns (middle); cell bodies in MGB/PIL region (right). (B) Experimental design to label pIns_PAG_ projecting neurons (left); axon terminals in PAG (middle); layer 5 cell bodies in pIns (right). (C) Experimental design to identify projections to vocal-gating region in the PAG (left); axon terminals of pIns_PAG_ projections (green) and cell bodies of PAG_RAm_ neurons (red); (D) Experimental design to identify pIns_PAG_ neurons that receive direct inputs from MGP_pIns_ neurons (left); layer 5 pIns_PAG_ neurons that expressed a colorflipper virus and switched from red to green due to the presence of Cre (right). Abbreviations: pIns, posterior insula; MGB, auditory thalamus; PIL, posterior intrathalamic nucleus; PAG, periaqueductal grey; RAm, nucleus retroambiguus.

Prior studies have established that neurons in the caudolateral PAG (clPAG) gate USV production in male mice through their axonal projections to vocal premotor neurons in the nucleus retroambiguus (PAG_RAm_)^24,37^. We explored whether pIns innervates this vocal gating region of the clPAG by injecting AAV retro-Cre in the PAG and AAV-FLEXed GFP in the pIns (Fig. 5B left, N = 4). This intersectional approach resulted in robust GFP expression in cell bodies in the pIns, especially in layer V (Fig. 5B, middle and right). We extended this approach by combining this intersectional with injections of retrogradely transported fluorescent latex microbeads into nucleus retroambiguus (RAm, Fig. 5C left). This approach revealed that the axon terminals of PAG-projecting pIns neurons overlapped with the region of clPAG that projects to RAm (Fig. 5C right, N = 2). Finally, we tested whether the region of the pIns that projects to PAG_RAm_ also receives input from the auditory thalamus (Fig. 5D left, N = 3). We injected AAV1-Cre into the MGB, resulting in Cre expression in pIns neurons postsynaptic to the MGB, and injected a retrogradely transported AAV into the clPAG, resulting in expression in the pIns of a fluorescent reporter that flips from red to green in a Cre-dependent manner (AAVrg-colorflipper). This approach resulted in GFP expression in cell bodies of layer 5 neurons in pIns (Fig. 5D right). These results indicate that the pIns directly links the auditory thalamus with the vocal-gating region in PAG.

### The posterior insula communicates with other brain regions for social behavior

We also performed further viral tracing experiments to map the efferents and afferents of the pIns. In one set of mice (N= 7 mice; 4 male and 3 female), we injected AAV9 in pIns of wildtype mice (N = 7) to express EGFP in pIns axonal efferents (Fig. 6A). In another set of mice (N= 7 mice; 4 male and 3 female), we injected AAVrg-Cre in pIns of transgenic Ai14 mice (N = 7) to express tdTomato in neurons afferent to the pIns (Fig 6B). These viral tracing experiments revealed strong reciprocal connections between the pIns and other cortical regions, including the anterior insula, amygdala, motor cortex, orbitofrontal cortex, piriform cortex and rhinal cortex (Fig. 6 C-F). Additionally, the pIns made reciprocal connections with the temporal association cortex, a region involved in encoding ultrasonic pup vocalizations^38^, and with the MGB/PIL. This approach also revealed that the pIns made a variety of non-reciprocal connections, especially with subcortical regions. These include efferents from the pIns to the PAG and afferents from the dorsal raphe nucleus and the ventromedial thalamic nucleus to the pIns. (Fig. 6C and E lower left). The current mapping results are consistent with an earlier study indicating that the pIns is bidirectionally connected with many cortical regions and mostly unidirectionally connected with subcortical connections^39^.

**Figure 6:**
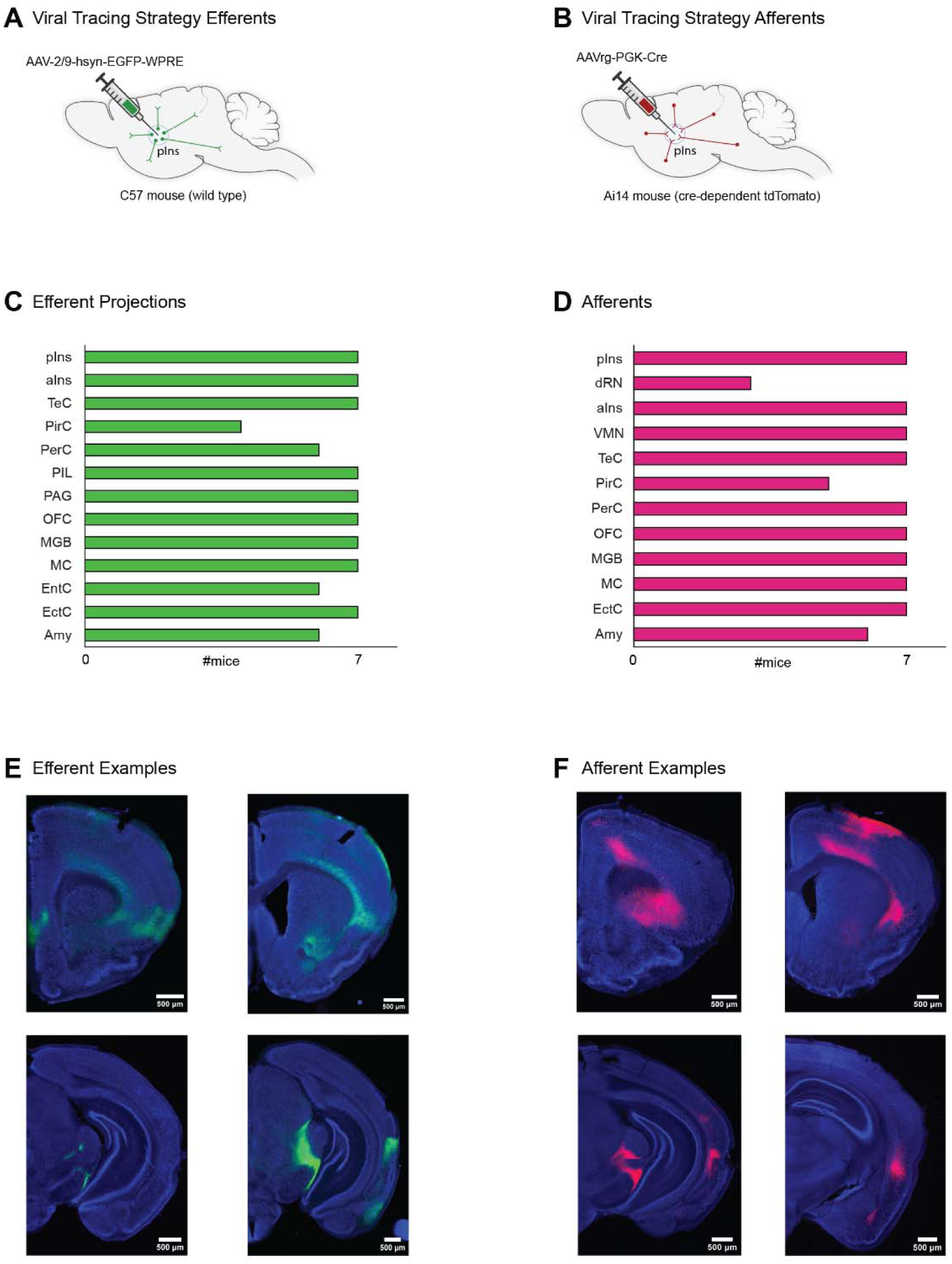
The posterior insula communicates with other brain regions for social behavior. (A-B) Experimental approach to trace efferents (green) and afferents (red) of the posterior insula. (C) Identified efferents in seven mice. (D) Identified afferents in seven mice. (E) Efferent examples. (F) Afferent examples. Abbreviations: pIns, posterior insula; aIns, anterior insula; TeC, temporal association cortex; PirC, piriform cortex; PerC, perirhinal cortex; PIL, posterior intrathalamic nucleus; PAG, periaqueductal grey; OFC, orbitofrontal cortex; MGB, medial geniculate body; MC, motor cortex; EntC, entorhinal cortex; EctC, ectorhinal cortex; Amy, amygdala; dRN, dorsal raphe nucleus; VMN, ventromedial thalamic nucleus;

## DISCUSSION

Here we used calcium imaging in freely courting male and female mice to characterize the activity of pIns neurons during vocal communication. In male mice, we identified two mostly distinct but spatially intermingled neuronal populations in the pIns that increased their activity during vocal communication. One population increased its activity prior to and during USV production in both hearing and deaf mice, consistent with a premotor or proprioceptive signal. Another population, which responded to USVs produced by nearby male mice engaged in courtship, was detected only in hearing and not deaf mice; similar responses were also detected in the pIns of female mice interacting with a vocalizing male. Notably, this USV-responsive population was only weakly excited by USV playback when mice were placed in social isolation, indicating that USV-responsiveness in the pIns is augmented by non-auditory social cues. In fact, 2-photon calcium imaging in head-fixed male mice revealed that female odorants could enhance USV responsiveness. A combination of intersectional and conventional tracing methods indicated that the pIns, and specifically layer 5 neurons, bridge the auditory thalamus with a region of the PAG that gates USV production. In summary, the pIns integrates auditory, olfactory and vocalization-related signals to encode expressive and receptive aspects of vocal communication in a manner that is sensitive to social context.

While the pIns is well known as a site for the auditory encoding of conspecific vocalizations^12^, pointing to a receptive function, here we found that the pIns is remarkably active during USV production, consistent with an expressive function. Specifically, activity in the pIns increased prior to the onset of USV production and remained elevated throughout the vocal bout. Moreover, similar patterns of vocalization-related activity were observed in hearing and congenitally deaf mice. The pre-vocal, hearing-independent nature of this vocalization-related activity is consistent with a motor-related signal. Indeed, a major afferent to the pIns identified here and in earlier studies^39^ is the secondary motor cortex (M2), a region that displays vocalization-related activity^40,41^ and that is a source of motor-related corollary discharge signals to the primary auditory cortex^28,29^. Another possibility is that the pIns integrates proprioceptive signals originating from respiratory and vocal muscles, although neither the somatosensory cortex or thalamus were labeled by the intersectional, retrograde tracing methods we employed. Because courtship USVs of male mice are typically produced in response to female odorants^20^, the pre-vocal activity in the pIns could also be linked to olfactory signals, which could be transmitted to the pIns from the piriform cortex and amygdala. However, delivering female odorants to a head-fixed male did not modulate pIns activity in the absence of subsequent USV production. A remaining possibility is that the pre-vocal signature in the pIns reflects signals related to the decision to vocalize, which could be transmitted to the pIns from the orbitofrontal cortex^42^ or from the anterior insula, the latter of which receives input from the medial prefrontal cortex^43^, a region directly linked to vocal production in rats, mice and monkeys^44–46^. In summary, a distinct subset of neurons the pIns are activated during vocal expression, mostly likely reflecting signals linked to vocal motor production or the decision to initiate vocalizations.

Previous studies found that the pIns responded to pure tones in mice^13,14^ and to vocalizations of conspecifics in rhesus monkeys^12^. While these studies point to the pIns as a site where vocalizations could be encoded in a socially salient manner, they monitored neural activity of playback stimuli delivered to head-fixed subjects in social isolation. An important advance of the current study is the analysis of pIns neurons during more naturalistic vocal communication involving several mice. Courtship USVs of male mice are typically produced in response to female odorants and render females more receptive to mating^20,23,24^. While male mice generate USVs without apparent intentionality or awareness as to outcome, the male’s USVs - given their pro-mating effects - can be regarded as adaptive signals that convey information about the male’s presence and reproductive fitness to an intended female target. However, as with other vocalizations, the courting male’s USVs convey information to any nearby animals that can hear them, including rival males. Here we created a social-vocal context in which a courting male’s USVs could be monitored by both a female mouse, the male’s intended courtship target, as well as a nearby “eavesdropping” male. This approach revealed that hearing the courting male’s USVs increased activity in neurons in the pIns in both the female and the eavesdropping male. This USV-evoked increase in activity was not simply a consequence of auditory stimulation, because USV playback evoked much less activity in the pIns when the female or the eavesdropping male was placed in social isolation. Instead, our results indicate that the pIns encodes male courtship USVs in a socially salient manner, consistent with prior studies that implicate the insula more generally in salience detection^5,47,48^.

The current study identifies female odorants as an important social cue that augments pIns activity in a male listening to another male’s USVs. Specifically, multiphoton imaging in head-fixed male mice revealed that exposure to female odorants augments responses in a male’s pIns to USV playback. Consistent with prior anatomical studies^6,39,49^, tracing experiments conducted here show that the pIns receives direct inputs from piriform cortex and amygdala, providing a pathway by which odorants could modulate USV responses. While we did not explore whether male odorants modulate USV-evoked responses in the pIns of the female mouse, these pathways are sexually monomorphic, suggesting that the pIns may serve a similar role in male and female mice. More broadly, odorants from mouse pups can modulate auditory cortical responses in dams to pup cries^50–52^, and thus odorant-dependent modulation may reflect a more general feature of the cortical representation of vocal sounds in the mouse cortex. Two-photon imaging in the pIns of socially isolated mice also revealed that a larger subset of neurons were modulated by USV playback than when the same region was imaged with 1p miniscopes in socially isolated unrestrained conditions. This could reflect the higher sensitivity of 2p methods or a heightened state of arousal in the head-fixed male that increases responses to auditory stimuli. Nevertheless, our results indicate that female odorants enhance responsiveness of pIns neurons in male mice listening to the courtship USVs of other males.

The present study also underscores the pivotal position of the pIns in the vocal sensorimotor hierarchy. The pIns is partly defined as a region that receives input from the auditory thalamus^15^. Our results extend these findings by elucidating that layer 5 neurons in the pIns receive direct input from the auditory thalamus and make axonal projections to USV-gating region in the PAG^24,37^. Whether these axons project directly to USV-gating neurons is unknown, but prior intersectional tracing studies from our lab suggest that they predominantly target local interneurons that provide inhibitory input onto USV-gating neurons in the PAG^24^. In this framework, activity evoked in the pIns by listening to another male’s USVs could serve to suppress USV production in the listener. However, a purely suppressive effect of the pIns on vocal gating neurons in the PAG cannot account for how activity in some pIns neurons increases before and during USV production. Therefore, an important goal of future studies will be to establish the identity, connectivity and function of pIns neurons that are active during expressive and receptive phases of social-vocal communication.

Effective social communication depends on establishing a correspondence between expressive and receptive aspects of communication signals. The current study shows that the pIns is a site where both expressive and receptive aspects of vocal signals are encoded, albeit in largely distinct neuronal populations. These observations confirm and extend a recent study in humans^16^ showing that the posterior insula is active during speech production and perception. Similar to primary auditory cortical neurons, we found that a population of pIns neurons that were responsive during vocal reception were suppressed during vocal production^16,30,53–57^. However, unlike the primary auditory cortex, an even larger subset of pIns neurons were strongly excited during vocal production, and this excitation arose earlier relative to vocal onset^30^. Therefore, the insula contains both expressive and receptive representations of vocal sounds, which could help to establish a sensorimotor correspondence that facilitates communication.

## MATERIALS AND METHODS

### Experimental models and subject details

#### Animals statement

All experiments were conducted according to a protocol approved by the Duke University Institutional Animal Care and Use Committee (protocol # A183-23-09 (1)).

#### Animals

For calcium imaging (1-photon and 2-photon) and neuronal tracing experiments, the following mouse lines from Jackson labs were used: C57 (C57BL/6J, Jackson Labs, 000664), Tmc1(Δ) (courtesy of Jeffery Holt, Harvard University) and Ai14 (B6.Cg-Gt(ROSA)26Sortim14(Cag-tdTomato)Hze/J, Jackson Labs, 007914). Mice were housed in 12/12 hours day/night cycle.

### Method details

#### Lens implantation and baseplating

One surface of a GRIN lense (4mm length, 1mm diameter, Inscopix) was covered with a silk-fibroin-virus mixture (1 part virus, 1 part silk fibroin) either the day before surgery and kept overnight at 4 degree Celsius or 30 minutes before implantation as described in (Jackman et al., 2018). Mice were then anesthetized (1.5%-2% isoflurane), and the pIns was targeted for injection. GRIN lenses were then implanted 0.1mm above pIns target location and were fixed to the skull using Metabond (Parkell) and dental cement (Ortho-Jet). We covered the lens with body-double and an additional layer of dental cement to protect it from damage. After a recovery period of 4-6 weeks, a baseplate was cemented on top of the animal and imaging experiments were conducted starting 3-7 days after baseplating.

#### USV recording and analysis

USVs were recorded using ultrasonic microphones (Avisoft, CMPA/CM16) amplified (Presonus TubePreV2), and digitized at 192 kHz/250 kHz (RZ6 Multi I/O Processor from Tucker Davis and a Power1401 CED board, Spike2) during 1p- and 2p-imaging, respectively. USVs were detected using Mupet^58^. USV bouts were defined by a minimum duration of 500ms and a minimum interbout duration of 2 seconds. Custom Matlab code was used to visualize each detected bout, and on- and offsets were manually adjusted if necessary.

#### Playback stimulus presentation

We used pre-recorded USVs from freely interacting males and females. Ultrasonic loud speakers (ES1 SN: 4907, Tucker Davis Technologies) were used to present these stimuli. Four different USV bouts with a length of 2-8 seconds were presented during the 1p-imaging experiments (10 presentations per stimulus, pseudorandomized order, 40 presentations in total). Six different USV bouts with a length of 2 seconds were presented during the 2p-imaging experiments (20 presentations per stimulus, pseudorandomized order, 120 presentations in total).

#### Behavior recording and analysis

All experiments were conducted under infrared light (IR Illuminator, model: YY-IR30W, LineMax). We used a webcam (HD 1080p, Logitech) from which we removed the infrared filter to monitor the behaviors of the mice. Animal pose estimations were acquired by using Deeplabcut^59^. We then used custom Matlab code to calculate speed and acceleration of an animal. Runnning bouts were defined as follows: minimum duration 0.5sec, interbout duration 1sec. Acceleration bouts were defined as follows: minimum duration 0.25sec. Area dimensions of the arena were acquired manually and we used video frames to convert pixels into metric values.

#### One-photon imaging

On the day of testing, a miniature miscroscope (UCLA miniscope V4) was mounted on the baseplate of the animal and fixated in place by a screw before the animal was placed into one of the two chambers of the two-chamber assay. Calcium data was acquired using the provided open-source software for UCLA miniscopes V4 which synchronized its recording times by sending out a TTL pulse to the audio recording system each time a frame was acquired. After an acclimation period of 3-5 minutes, the animal was exposed to other conspecifics and playback stimuli. Video, audio and calcium signals were recorded as the mouse freely interacted with the presented stimuli. The resulting calcium signal was analyzed using Minian^60^ and custom Matlab codes. Extracted ROIs were manually inspected.

#### Two-photon imaging

Prior to 2-photon calcium imaging we implanted titanium Y-headbars on mice using Metabond (Parkell) after they underwent surgery for GRIN lens implantation as described above. Mice were head-fixed on a radial treadmill and habituated for at least one week before the experiment was conducted. The baseplate was filled with carbomer gel (refractive index 1.4) and signal were recorded by a 10x/0.45NA water immersion objective (Nikon). We used a titanium sapphire laser (MaiTai DeepSee, 920nm, Neurolabware) with a laser power of 100mW. Recordings were performed in darkness. Data was acquired using Scanbox (sampling rate 15.49 Hz; 512 x 512 pixels) that sent out a TTL pulse to Spike7 audio-recording system each time a frame was acquired. Suite2p^61^ was used to extract individual calcium signals and subsequent data analysis was performed by custom Matlab code. Extracted ROIs were manually inspected.

#### Viruses and tracers

We used the following viruses and tracers: AAV2/9-syn-jGCaMP8s-WPRE (Addgene), AAVrg-PGK-Cre (Addgene), AAV-2/1/CAG-Flex-EGFP (Addgene), pENN.AAV.hsyn.Cre.WPRE.hGH (AAV1, Addgene), pOOTC1032 – pAAV-EF1a-Nuc-flox(mCherry)-EGFP and Red Retrobeads™ IX (LumaFluor). We injected into the following coordinates relative to bregma: pIns, AP=-1.05mm, ML=3.80mm, DV=-3.50mm; MGB, AP=-2.90mm, ML=-1.75mm, DV=-3.40mm; PAG=-4.7mm, ML=0.70mm, DV=-1.75mm; RAm: AP=-8.05mm, ML=1.00mm, DV=-5.20mm. Coordinates were achieved via a digital stereotaxic instrument (RWD) and viruses were pressure-injected with a Nanoject III (Drummond) at a rate of 1nl/sec.

#### Post-hoc visualization of viral labeling

Mice were deeply anesthetized with isoflurance and then transcardially perfused with ice-cold 4% paraformaldehyde in 0.1 M phosphate buffer, pH 7.4 (4% PFA). Dissected brain samples were postfixed overnight in 4% PFA at 4 degrees C, cryoprotected in a 30% sucrose solution in PBS at 4 degrees C for 48 hrs, frozen in Tissue-Tek O.C.T. Compound (Sakura), and stored at - 80 degrees Celcius until sectioning. Brains were cut into 100 μm coronal sections, rinsed 3x in PBS, and processed for 24 hrs at 4 degrees with NeuroTrace (1:500, Invitrogen) in PBS. To increase fluorescence of jGCaMP8s in brain slices we added primary antibody (Chk pAb to GFP, ab13970, Abcam) to NeuroTraces, rinsed the samples 3x in PBS and processed with secondary antibody (Anti-Chicken IgY, 488, 703-545-155, Jackson ImmunoResearch). Tissue sections were rinsed again 3x in PBS, mounted on slides, and coverslipped with Fluoromount-G (Southern Biotech). After drying, slides were imaged with a 10x objective in a Zeiss 700 laser scanning confocal microscope and a Keyence microscope (BZ-X810, All-in-One Fluorescence Microscope).

#### Statistical analysis

All ΔF/F calcium traces were z-scored prior to each analysis and were presented in units of standard deviation. We quantified responses of each neurons during miniscope recordings using a receiver operating characteristic (ROC) analysis, which has been applied previously to detect responses during natural behavior^62,63^. We calculated ROC curves for each ROI by first obtaining the distribution of calcium responses across all vocal bouts and during baseline (difference of means before and after vocal onset) and then used a moving criterion from minimum calcium amplitude to maximum calcium amplitude of those two distributions. The length of this moving criterion was calculated as the (max – min)/100. We then used the area under the resulting ROC curve and compared it to a 1000 times randomly shuffled distribution for each ROI. ROIs with ROC area under the curve values below or above the 2.5^th^ and 97.5^th^ percentile were considered suppressed or excited during vocal bouts, respectively. Playback responses during 1-photon and 2-photon calcium imaging and changes in running speed were quantified using a two-sided Wilcoxon ranksum test. Individual average calcium signals were baseline subtracted prior to visualization.

#### Decoding analysis

For each recording session we performed a principal component analysis on all ROIs and used the first 24 principle components (PCs) as the input layer to our model. We then divided each recording session into two data sets: The training data set contained 85% of vocal bouts while the test data set contained 15% of vocal bouts. We then created a long short-term memory network using Matlab that we trained on the training data set to decode vocal bouts by PC activity. Next, we applied the decoder to the original test data and a control data set where we randomly shuffled the USV bout appearances. The resulting decoding accuracies for each session were quantified by a standard two-sided Wilcoxon ranksum test.

### Aknowledgements

The authors would like to thank Professor Jeffery Holt (Harvard Medical School) for donating *Tmc1*Δ/Δ mice and Michael Booze for animal husbandry and genotyping. They also thank all members of the Mooney lab for their helpful discussion and support. This research was supported by grants from the National Institutes of Health: R01DC013826-07 (R.M.), and R01MH117778-05 (R.M.).

---

**Supplemental figure 1:**
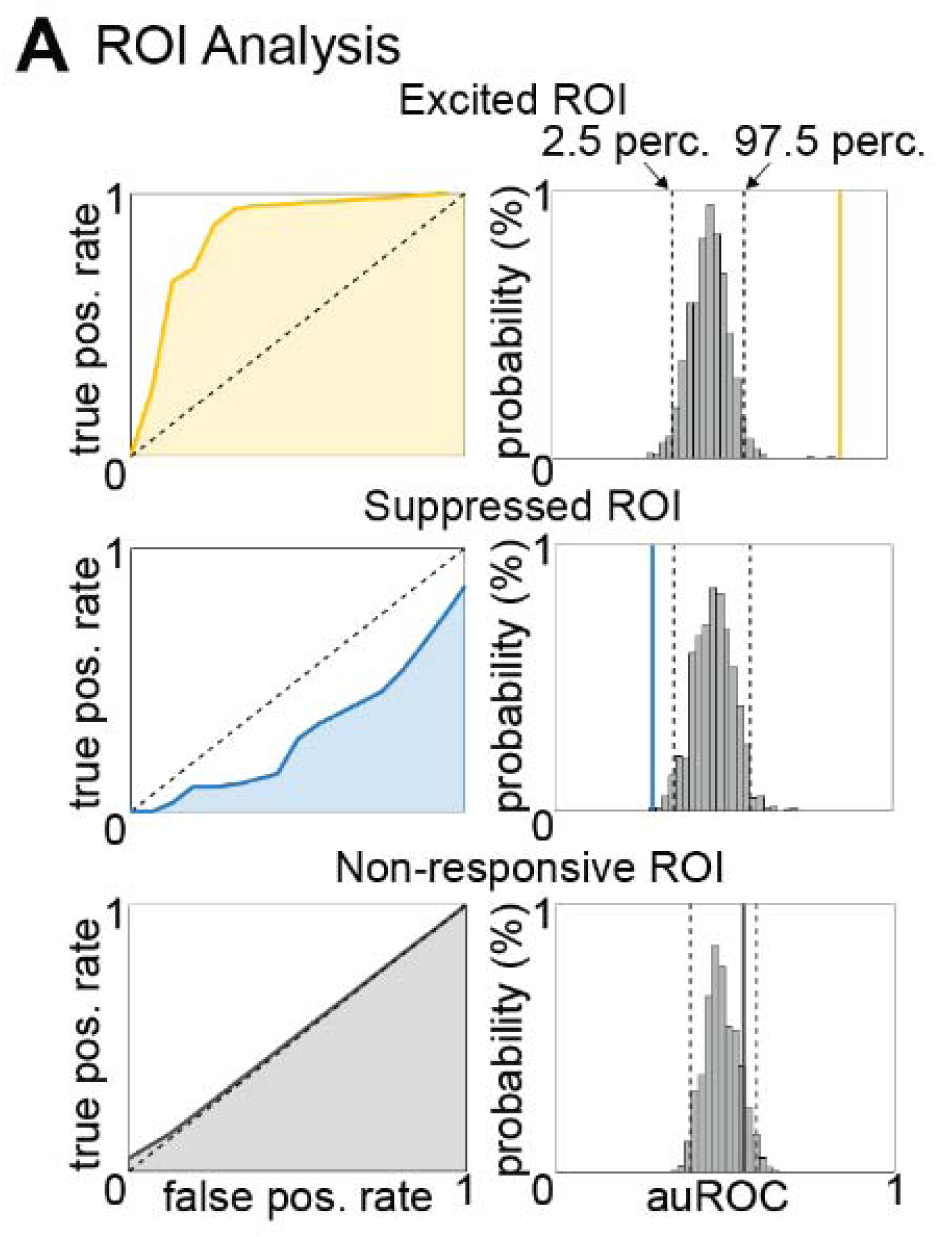
(A) Schematic examples of receiver-operator characteristic to quantif responsiveness of ROIs: top, excited; middle, suppressed; bottom, non-responsive.

**Supplemental figure 2:**
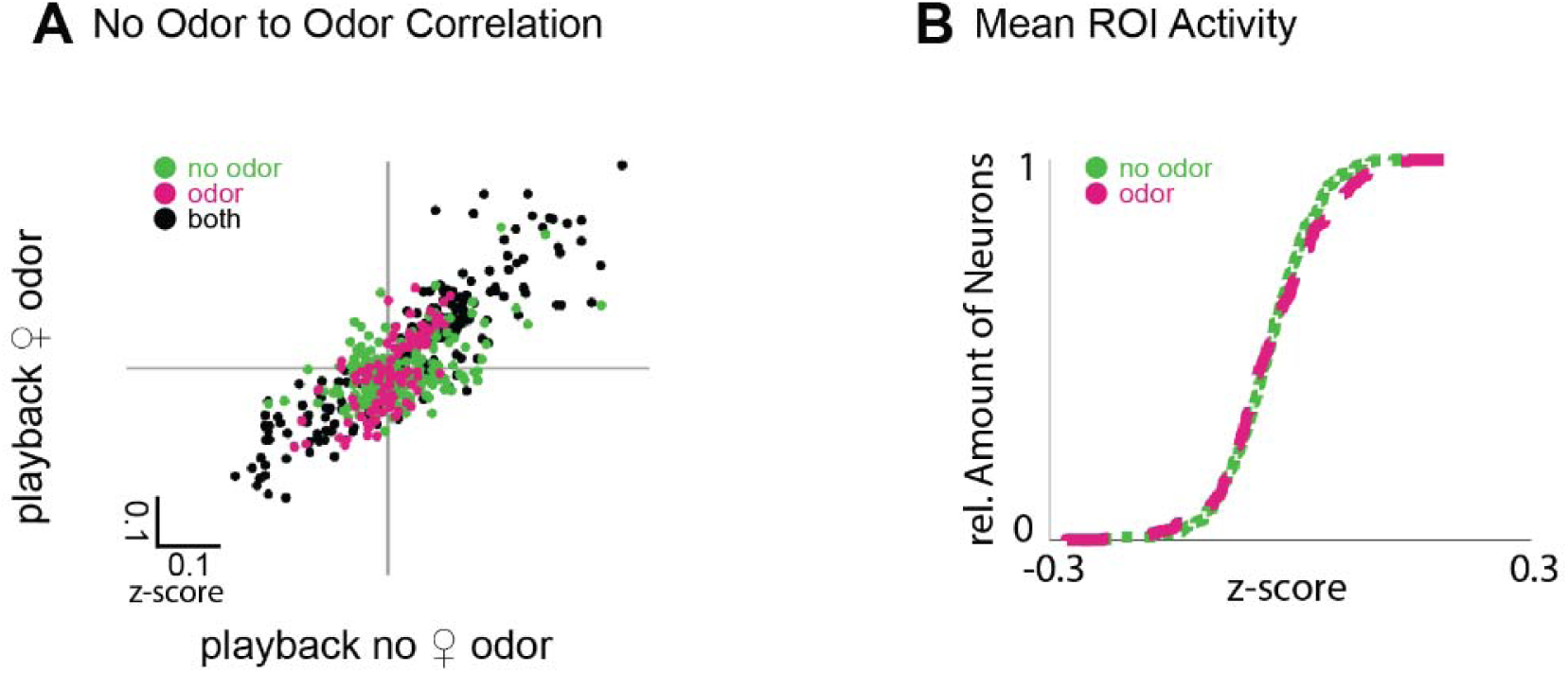
(A) Correlation of mean activities of ROIs responsive during USV playback. Colors depict responsiveness in neutral airflow (green), positive airflow (magenta) and in both (black). (B) Cumulative distribution function of baseline activity during neutral airflow (green) and positive airflow (magenta).

## References

1. Wright, G.S., Chiu, C., Xian, W., Wilkinson, G.S., and Moss, C.F. (2013). Social calls of flying big brown bats (Eptesicus fuscus). Front. Physiol. 4. 10.3389/fphys.2013.00214.

2. Williams, J.H.G., Huggins, C.F., Zupan, B., Willis, M., Van Rheenen, T.E., Sato, W., Palermo, R., Ortner, C., Krippl, M., Kret, M., et al. (2020). A sensorimotor control framework for understanding emotional communication and regulation. Neuroscience & Biobehavioral Reviews 112, 503–518. 10.1016/j.neubiorev.2020.02.014.

3. Chereskin, E., Allen, S.J., Connor, R.C., Krützen, M., and King, S.L. (2024). In pop pursuit: social bond strength predicts vocal synchrony during cooperative mate guarding in bottlenose dolphins. Phil. Trans. R. Soc. B 379, 20230194. 10.1098/rstb.2023.0194.

4. Warren, M.R., Young, L.J., and Liu, R.C. (2024). Vocal recognition of partners by female prairie voles. Preprint, 10.1101/2024.07.24.604991.

5. Uddin, L.Q. (2015). Salience processing and insular cortical function and dysfunction. Nat Rev Neurosci 16, 55–61. 10.1038/nrn3857.

6. Gogolla, N. (2017). The insular cortex. Current Biology 27, R580–R586. 10.1016/j.cub.2017.05.010.

7. Livneh, Y., Ramesh, R.N., Burgess, C.R., Levandowski, K.M., Madara, J.C., Fenselau, H., Goldey, G.J., Diaz, V.E., Jikomes, N., Resch, J.M., et al. (2017). Homeostatic circuits selectively gate food cue responses in insular cortex. Nature 546, 611–616. 10.1038/nature22375.

8. Livneh, Y., Sugden, A.U., Madara, J.C., Essner, R.A., Flores, V.I., Sugden, L.A., Resch, J.M., Lowell, B.B., and Andermann, M.L. (2020). Estimation of Current and Future Physiological States in Insular Cortex. Neuron 105, 1094–1111.e10. 10.1016/j.neuron.2019.12.027.

9. Dolensek, N., Gehrlach, D.A., Klein, A.S., and Gogolla, N. (2020). Facial expressions of emotion states and their neuronal correlates in mice. Science 368, 89–94. 10.1126/science.aaz9468.

10. Klein, A.S., Dolensek, N., Weiand, C., and Gogolla, N. (2021). Fear balance is maintained by bodily feedback to the insular cortex in mice. Science 374, 1010–1015. 10.1126/science.abj8817.

11. Livneh, Y., and Andermann, M.L. (2021). Cellular activity in insular cortex across seconds to hours: Sensations and predictions of bodily states. Neuron 109, 3576–3593. 10.1016/j.neuron.2021.08.036.

12. Remedios, R., Logothetis, N.K., and Kayser, C. (2009). An Auditory Region in the Primate Insular Cortex Responding Preferentially to Vocal Communication Sounds. J. Neurosci. 29, 1034–1045. 10.1523/JNEUROSCI.4089-08.2009.

13. Sawatari, H., Tanaka, Y., Takemoto, M., Nishimura, M., Hasegawa, K., Saitoh, K., and Song, W. (2011). Identification and characterization of an insular auditory field in mice. Eur J of Neuroscience 34, 1944–1952. 10.1111/j.1460-9568.2011.07926.x.

14. Gogolla, N., Takesian, A.E., Feng, G., Fagiolini, M., and Hensch, T.K. (2014). Sensory Integration in Mouse Insular Cortex Reflects GABA Circuit Maturation. Neuron 83, 894–905. 10.1016/j.neuron.2014.06.033.

15. Takemoto, M., Hasegawa, K., Nishimura, M., and Song, W. (2014). The insular auditory field receives input from the lemniscal subdivision of the auditory thalamus in mice. J of Comparative Neurology 522, 1373–1389. 10.1002/cne.23491.

16. Kurteff, G.L., Field, A.M., Asghar, S., Tyler-Kabara, E.C., Clarke, D., Weiner, H.L., Anderson, A.E., Watrous, A.J., Buchanan, R.J., Modur, P.N., et al. (2024). Processing of auditory feedback in perisylvian and insular cortex. Preprint, 10.1101/2024.05.14.593257.

17. Dronkers, N.F. (1996). A new brain region for coordinating speech articulation. Nature 384, 159–161. 10.1038/384159a0.

18. Dronkers, N.F., Plaisant, O., Iba-Zizen, M.T., and Cabanis, E.A. (2007). Paul Broca’s historic cases: high resolution MR imaging of the brains of Leborgne and Lelong. Brain 130, 1432–1441. 10.1093/brain/awm042.

19. Sewell, G.D.S.N. (1972). Ultrasound and mating behaviour in rodents with some observations on other behavioural situations. Journal of Zoology 168, 149–164. 10.1111/j.1469-7998.1972.tb01345.x.

20. Whitney, G., Alpern, M., Dizinno, G., and Horowitz, G. (1974). Female odors evoke ultrasounds from male mice. Animal Learning & Behavior 2, 13–18. 10.3758/BF03199109.

21. Dizinno, G., Whitney, G., and Nyby, J. (1978). Ultrasonic vocalizations by male mice (Mus musculus) to female sex pheromone: Experiential determinants. Behavioral Biology 22, 104–113. 10.1016/S0091-6773(78)92094-1.

22. Portfors, C.V., and Perkel, D.J. (2014). The role of ultrasonic vocalizations in mouse communication. Current Opinion in Neurobiology 28, 115–120. 10.1016/j.conb.2014.07.002.

23. Pomerantz, S.M., Nunez, A.A., and Jay Bean, N. (1983). Female behavior is affected by male ultrasonic vocalizations in house mice. Physiology & Behavior 31, 91–96. 10.1016/0031-9384(83)90101-4.

24. Tschida, K., Michael, V., Takatoh, J., Han, B.-X., Zhao, S., Sakurai, K., Mooney, R., and Wang, F. (2019). A Specialized Neural Circuit Gates Social Vocalizations in the Mouse. Neuron 103, 459–472.e4. 10.1016/j.neuron.2019.05.025.

25. Neunuebel, J.P., Taylor, A.L., Arthur, B.J., and Egnor, S.R. (2015). Female mice ultrasonically interact with males during courtship displays. eLife 4, e06203. 10.7554/eLife.06203.

26. Sterling, M.L., Teunisse, R., and Englitz, B. (2023). Rodent ultrasonic vocal interaction resolved with millimeter precision using hybrid beamforming. eLife 12, e86126. 10.7554/eLife.86126.

27. Waidmann, E.N., Yang, V.H.Y., Doyle, W.C., and Jarvis, E.D. (2024). Mountable miniature microphones to identify and assign mouse ultrasonic vocalizations. Preprint, 10.1101/2024.02.05.579003.

28. Schneider, D.M., Nelson, A., and Mooney, R. (2014). A synaptic and circuit basis for corollary discharge in the auditory cortex. Nature 513, 189–194. 10.1038/nature13724.

29. Schneider, D.M., Sundararajan, J., and Mooney, R. (2018). A cortical filter that learns to suppress the acoustic consequences of movement. Nature 561, 391–395. 10.1038/s41586-018-0520-5.

30. Harmon, T.C., Madlon-Kay, S., Pearson, J., and Mooney, R. (2024). Vocalization modulates the mouse auditory cortex even in the absence of hearing. Cell Reports 43, 114611. 10.1016/j.celrep.2024.114611.

31. Kawashima, Y., Géléoc, G.S.G., Kurima, K., Labay, V., Lelli, A., Asai, Y., Makishima, T., Wu, D.K., Della Santina, C.C., Holt, J.R., et al. (2011). Mechanotransduction in mouse inner ear hair cells requires transmembrane channel–like genes. J. Clin. Invest. 121, 4796–4809. 10.1172/JCI60405.

32. Rodgers, K.M., Benison, A.M., Klein, A., and Barth, D.S. (2008). Auditory, Somatosensory, and Multisensory Insular Cortex in the Rat. Cerebral Cortex 18, 2941–2951. 10.1093/cercor/bhn054.

33. Hansen, S., and Köhler, C. (1984). The Importance of the Peripeduncular Nucleus in the Neuroendocrine Control of Sexual Behavior and Milk Ejection in the Rat. Neuroendocrinology 39, 563–572. 10.1159/000124038.

34. Dobolyi, A., Cservenák, M., and Young, L.J. (2018). Thalamic integration of social stimuli regulating parental behavior and the oxytocin system. Frontiers in Neuroendocrinology 51, 102–115. 10.1016/j.yfrne.2018.05.002.

35. Valtcheva, S., Issa, H.A., Bair-Marshall, C.J., Martin, K.A., Jung, K., Zhang, Y., Kwon, H.-B., and Froemke, R.C. (2023). Neural circuitry for maternal oxytocin release induced by infant cries. Nature 621, 788–795. 10.1038/s41586-023-06540-4.

36. Leithead, A.B., Godino, A., Barbier, M., and Harony-Nicolas, H. (2024). Social Interaction Elicits Activity in Glutamatergic Neurons in the Posterior Intralaminar Complex of the Thalamus. Preprint, 10.1101/2023.04.24.538114.

37. Michael, V., Goffinet, J., Pearson, J., Wang, F., Tschida, K., and Mooney, R. (2020). Circuit and synaptic organization of forebrain-to-midbrain pathways that promote and suppress vocalization. eLife 9, e63493. 10.7554/eLife.63493.

38. Tasaka, G., Feigin, L., Maor, I., Groysman, M., DeNardo, L.A., Schiavo, J.K., Froemke, R.C., Luo, L., and Mizrahi, A. (2020). The Temporal Association Cortex Plays a Key Role in Auditory-Driven Maternal Plasticity. Neuron 107, 566–579.e7. 10.1016/j.neuron.2020.05.004.

39. Gehrlach, D.A., Weiand, C., Gaitanos, T.N., Cho, E., Klein, A.S., Hennrich, A.A., Conzelmann, K.-K., and Gogolla, N. (2020). A whole-brain connectivity map of mouse insular cortex. eLife 9, e55585. 10.7554/eLife.55585.

40. Okobi, D.E., Banerjee, A., Matheson, A.M.M., Phelps, S.M., and Long, M.A. (2019). Motor cortical control of vocal interaction in neotropical singing mice. Science 363, 983–988. 10.1126/science.aau9480.

41. Banerjee, A., Chen, F., Druckmann, S., and Long, M.A. (2024). Temporal scaling of motor cortical dynamics reveals hierarchical control of vocal production. Nat Neurosci 27, 527–535. 10.1038/s41593-023-01556-5.

42. Wallis, J.D. (2007). Orbitofrontal Cortex and Its Contribution to Decision-Making. Annu. Rev. Neurosci. 30, 31–56. 10.1146/annurev.neuro.30.051606.094334.

43. Gabbott, P.L.A., Warner, T.A., Jays, P.R.L., and Bacon, S.J. (2003). Areal and synaptic interconnectivity of prelimbic (area 32), infralimbic (area 25) and insular cortices in the rat. Brain Research 993, 59–71. 10.1016/j.brainres.2003.08.056.

44. Hage, S.R., and Nieder, A. (2013). Single neurons in monkey prefrontal cortex encode volitional initiation of vocalizations. Nat Commun 4, 2409. 10.1038/ncomms3409.

45. Bennett, P.J.G., Maier, E., and Brecht, M. (2019). Involvement of rat posterior prelimbic and cingulate area 2 in vocalization control. Eur J Neurosci 50, 3164–3180. 10.1111/ejn.14477.

46. Gan-Or, B., and London, M. (2023). Cortical circuits modulate mouse social vocalizations. Sci. Adv. 9, eade6992. 10.1126/sciadv.ade6992.

47. Crottaz-Herbette, S., and Menon, V. (2006). Where and When the Anterior Cingulate Cortex Modulates Attentional Response: Combined fMRI and ERP Evidence. Journal of Cognitive Neuroscience 18, 766–780. 10.1162/jocn.2006.18.5.766.

48. Bonnelle, V., Ham, T.E., Leech, R., Kinnunen, K.M., Mehta, M.A., Greenwood, R.J., and Sharp, D.J. (2012). Salience network integrity predicts default mode network function after traumatic brain injury. Proc. Natl. Acad. Sci. U.S.A. 109, 4690–4695. 10.1073/pnas.1113455109.

49. Ghaziri, J., Tucholka, A., Girard, G., Boucher, O., Houde, J.-C., Descoteaux, M., Obaid, S., Gilbert, G., Rouleau, I., and Nguyen, D.K. (2018). Subcortical structural connectivity of insular subregions. Sci Rep 8, 8596. 10.1038/s41598-018-26995-0.

50. Cohen, L., Rothschild, G., and Mizrahi, A. (2011). Multisensory Integration of Natural Odors and Sounds in the Auditory Cortex. Neuron 72, 357–369. 10.1016/j.neuron.2011.08.019.

51. Cohen, L., and Mizrahi, A. (2015). Plasticity during Motherhood: Changes in Excitatory and Inhibitory Layer 2/3 Neurons in Auditory Cortex. J. Neurosci. 35, 1806–1815. 10.1523/JNEUROSCI.1786-14.2015.

52. Gilday, O.D., and Mizrahi, A. (2023). Learning-Induced Odor Modulation of Neuronal Activity in Auditory Cortex. J. Neurosci. 43, 1375–1386. 10.1523/JNEUROSCI.1398-22.2022.

53. Creutzfeldt, O., Ojemann, G., and Lettich, E. (1989). Neuronal activity in the human lateral temporal lobe: II. Responses to the subjects own voice. Exp Brain Res 77, 476–489. 10.1007/BF00249601.

54. Eliades, S.J., and Wang, X. (2008). Neural substrates of vocalization feedback monitoring in primate auditory cortex. Nature 453, 1102–1106. 10.1038/nature06910.

55. Towle, V.L., Yoon, H.-A., Castelle, M., Edgar, J.C., Biassou, N.M., Frim, D.M., Spire, J.-P., and Kohrman, M.H. (2008). ECoG gamma activity during a language task: differentiating expressive and receptive speech areas. Brain 131, 2013–2027. 10.1093/brain/awn147.

56. Eliades, S.J., and Tsunada, J. (2018). Auditory cortical activity drives feedback-dependent vocal control in marmosets. Nat Commun 9, 2540. 10.1038/s41467-018-04961-8.

57. Tsunada, J., Wang, X., and Eliades, S.J. (2024). Multiple processes of vocal sensory-motor interaction in primate auditory cortex. Nat Commun 15, 3093. 10.1038/s41467-024-47510-2.

58. Van Segbroeck, M., Knoll, A.T., Levitt, P., and Narayanan, S. (2017). MUPET—Mouse Ultrasonic Profile ExTraction: A Signal Processing Tool for Rapid and Unsupervised Analysis of Ultrasonic Vocalizations. Neuron 94, 465–485.e5. 10.1016/j.neuron.2017.04.005.

59. Lauer, J., Zhou, M., Ye, S., Menegas, W., Schneider, S., Nath, T., Rahman, M.M., Di Santo, V., Soberanes, D., Feng, G., et al. (2022). Multi-animal pose estimation, identification and tracking with DeepLabCut. Nat Methods 19, 496–504. 10.1038/s41592-022-01443-0.

60. Dong, Z., Mau, W., Feng, Y., Pennington, Z.T., Chen, L., Zaki, Y., Rajan, K., Shuman, T., Aharoni, D., and Cai, D.J. (2022). Minian, an open-source miniscope analysis pipeline. eLife 11, e70661. 10.7554/eLife.70661.

61. Pachitariu, M., Stringer, C., Dipoppa, M., Schröder, S., Rossi, L.F., Dalgleish, H., Carandini, M., and Harris, K.D. (2016). Suite2p: beyond 10,000 neurons with standard two-photon microscopy. Preprint, 10.1101/061507.

62. Li, Y., Mathis, A., Grewe, B.F., Osterhout, J.A., Ahanonu, B., Schnitzer, M.J., Murthy, V.N., and Dulac, C. (2017). Neuronal Representation of Social Information in the Medial Amygdala of Awake Behaving Mice. Cell 171, 1176–1190.e17. 10.1016/j.cell.2017.10.015.

63. Kingsbury, L., Huang, S., Wang, J., Gu, K., Golshani, P., Wu, Y.E., and Hong, W. (2019). Correlated Neural Activity and Encoding of Behavior across Brains of Socially Interacting Animals. Cell 178, 429–446.e16. 10.1016/j.cell.2019.05.022.

